# LncRNA-BERT: An RNA Language Model for Classifying Coding and Long Non-Coding RNA

**DOI:** 10.1101/2025.01.09.632168

**Authors:** Luuk Romeijn, Davy Cats, Katherine Wolstencroft, Hailiang Mei

## Abstract

Understanding (novel) RNA transcripts generated in next generation sequencing experiments requires accurate classification, given the increasing evidence that long non-coding RNAs (lncRNAs) play crucial regulatory roles. Recent developments in Large Language Models present opportunities for classifying RNA coding potential with sequence-based algorithms that can overcome the limitations of classical approaches that assess coding potential based on a set of predefined features. We present lncRNA-BERT, an RNA language model pre-trained and fine-tuned on human RNAs collected from the GENCODE, RefSeq, and NONCODE databases to classify lncRNAs. LncRNA-BERT matches and outperforms state-of-the-art classifiers on three test datasets, including the cross-species RNAChallenge benchmark. The pre-trained lncRNA-BERT model distinguishes coding from long non-coding RNA without supervised learning which confirms that coding potential is a sequence-intrinsic characteristic. LncRNA-BERT has been shown to benefit from pre-training on human data from GENCODE, RefSeq, and NONCODE, improving upon configurations pre-trained on the commonly used RNAcentral dataset. In addition, we propose a novel Convolutional Sequence Encoding method that is shown to be more effective and efficient than K-mer Tokenization and Byte Pair Encoding for training with long RNA sequences that are otherwise above the common context window size. lncRNA-BERT is available at https://github.com/luukromeijn/lncRNA-Py.

## 1 Introduction

Owing to advancements in RNA library extraction and sequencing technologies, bulk RNA and single cell RNA sequencing have been regularly applied in research projects and routine clinical diagnostics. Bioinformatics analysis of these RNAseq datasets leverages the well-maintained annotations from gene annotation databases, such as RefSeq and Ensembl, to examine the expression profiles of genes or transcripts. While earlier research has focused on protein coding transcripts, more recent work has investigated non-coding RNAs (ncRNAs) for their important regulatory functions [1–3]. Non-coding RNAs are often classified into two large groups based on their transcript length: short ncRNAs (including biotypes such as miRNA, piRNA, snoRNA, and tRNA) and long non-coding RNAs (lncRNAs) with a length ≥ 200 nt. Specific databases, such as NONCODE and RNAcentral, have been implemented to track newly reported ncRNAs across different species [4, 5]. In fact, lncRNAs are a highly prevalent type of RNA, underlined by the 173,112 human lncRNA transcripts stored in the NONCODE (v6) database versus the 197,151 mRNAs in RefSeq (r225).

To maintain high-quality curation, it is important to classify novel transcripts into a proper biotype and, in particular, distinguish lncRNAs from protein-coding RNAs (pcRNA) as they often share a similar transcript length. Currently, researchers can choose between more than 40 algorithms to classify whether a novel transcript sequence is a lncRNA.

Most of these classifiers are machine learning models that have been trained on annotated RNA data from RefSeq or GENCODE (e.g. CPC, CNCI, CPAT, and CPC2 [6–9]), extracting features such as ORF length, protein database alignment hits, and k-mer frequencies as predictors for coding potential. The challenge of the classification problem lies in its inherent ambiguity: some lncRNAs contain short ORFs that translate into small peptide chains [10], and pcRNA genes can have non-coding isoforms [11,12]. This has stimulated the continuous development of novel classifiers, leveraging the latest available annotations and methodological advancements, such as feature selection algorithms [13,14], boosting models [15], and neural networks [16–18].

Although feature-based algorithms can learn the relationship between the target and a predetermined set of predictors, they may fail to capture the true underlying signal. Purely sequence-based deep learning methods for lncRNA classification have been proposed [17, 19, 20], but these were outperformed by feature-based or hybrid methods in a recent benchmark [21]. These approaches utilize convolutional and/or recurrent neural networks (CNN/RNN), which suffer from limited receptive fields and exploding/vanishing gradients, respectively. The transformer architecture improves upon these designs by employing an attention mechanism and is nonrecurrent [22].

We propose using a transformer-based Nucleotide Language Model (NLM) to overcome the limitations of previous classifiers. NLMs are Large Language Models (LLMs) that have been pre-trained and finetuned on genomic data. DNA NLMs have demonstrated their potential to predict chromatin features, identify promoter regions, and perform variant prioritization [23–25]. RNA NLMs have been shown to be capable of predicting splice sites, secondary structures, and interactions of RNA [26–30]. Despite being presented for general purposes, previous RNA language models may not be properly adapted for detecting coding potential due to utilizing RNAcentral (which excludes pcRNA) as the main data source and having a limited context length (*<* 1024 nt).

In this work, we present a new lncRNA classification method called lncRNA-BERT (long non-coding RNA Bidirectional Encoder Representations from Transfomers), an RNA language model specifically adapted to classifying RNA as coding or long non-coding. The main contributions of our study are threefold. First, lncRNA-BERT obtained state-of-the-art classification performance on three different test sets and significantly outperformed all other methods in the most difficult cross-species RNAChallenge test set. Second, lncRNA-BERT accurately distinguished between pcRNA and lncRNA without access to target labels by pre-training on human RNA from GENCODE, RefSeq, and NONCODE, rather than multi-species ncRNA from RNAcentral. Third, an in-depth comparison of four sequence encoding methods demonstrates the effectiveness of a novel Convolutional Sequence Encoding (CSE) method for pre-training on long RNA sequences and identifies CSE and 3-mer tokenization as the most suitable methods for classifying lncRNAs.

## 2 Methods

lncRNA-BERT was implemented in the Python package lncRNA-Py, which is available from GitHub (https://github.com/luukromeijn/lncRNA-Py). Additional experiments, such as architecture and hyperparameter tuning, have also been reported.

### 2.1 Data

An overview of the datasets utilized for the different tasks is presented in Table 1. The main pre-training dataset is a human set of 297,724 coding and 238,470 non-coding RNA sequences from GENCODE (v46) [31], NONCODE (v6) [4], and RefSeq (v255) [32]. A randomly selected 5% of the GENCODE sequences was held out for validation. We experimented with RNAcentral (v24) [5], which contains 37 million ncRNAs, as an alternative multi-species pre-training data source and kept aside 2,500 sequences for validating this model.

**Table 1:**
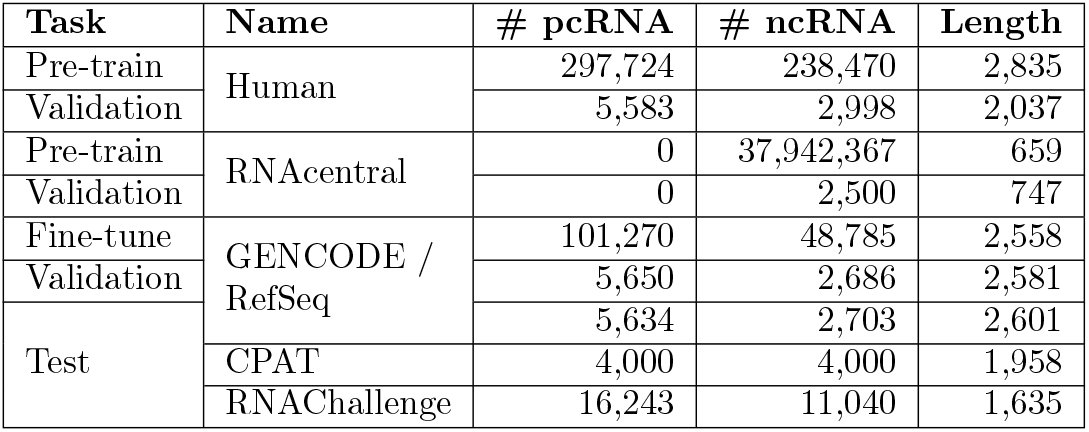
Data specification. Human data from GENCODE, RefSeq, and NONCODE is used. The three GENCODE/RefSeq datasets are obtained by first clustering the data with CD-HIT (90% identity threshold), and then randomly selecting 90%, 5%, and 5% for fine-tuning, validation, and testing, respectively. Average length is reported.

For fine-tuning, we used the CD-HIT algorithm [33] to ensure non-redundancy and test set independence, similar to [13, 15]. We ran the CD-HIT algorithm with a 90% sequence identity threshold on the combined human pcRNA/lncRNA data from GENCODE and RefSeq and randomly selected 90% of representative sequences for training, 5% for validation, and 5% for testing. The fine-tuned set contains 101,270 protein-coding and 48,785 non-coding RNAs. NONCODE was deliberately excluded during fine-tuning to maximize the data reliability.

Three test sets were used to assess the performance of lncRNA-BERT and the other classifiers: GENCODE/RefSeq, CPAT, and RNAChallenge. The GENCODE/RefSeq test set (5,650 pcRNAs, 2,686 lncRNAs) is guaranteed to not overlap with our fine-tuning data because of the above-described redundancy removal with CD-HIT. The CPAT set (4,000 pcRNAs, 4,000 lncRNAs) is a published and widely used test set [8], although some overlap with the training sets of each of the evaluated classifiers is expected. Finally, we used another published benchmark, RNAChallenge [21], containing 27,283 hard-to-classify RNA sequences, to assess the generalizability of our model to ambiguous RNA sequences from animal, plant, and fungi species.

### 2.2 Encoding Methods

We compared four methods for converting nucleotide sequences into a numerical format: Nucleotide-Level Tokenization (NUC), K-mer Tokenization (K-mer), Byte Pair Encoding (BPE), and a novel Convolutional Sequence Encoding (CSE) method. Efficient encoding is required for long sequences, as the transformer’s attention mechanism only accepts a limited number of input positions (512-1024 on standard GPUs) due to its quadratic memory complexity. The effects of different encoding methods on the encoded sequence length are illustrated in Fig.1.

**Figure 1:**
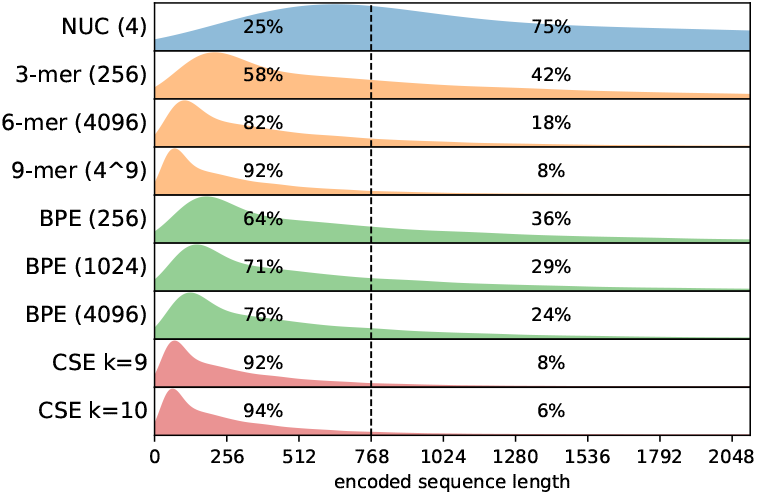
Density plots of encoded sequence lengths from the human pre-training dataset (536,194 sequences) when using NUC, K-mer Tokenization, BPE, and CSE. Vocabulary sizes are provided in parentheses. The percentages represent the number of encoded sequences that fall within a medium-sized context length of 768 input positions.

#### 2.2.1 Nucleotide-Level Tokenization

Most RNA NLMs apply NUC as the sequence encoding method using a vocabulary of four nucleotide tokens (A,C,T,G). This technique allows attention to nucleotide resolution and works well for short sequences, such as most RNAs in RNAcentral. However, NUC is insufficient for long RNAs in our human pre-training set, as only 25% would fit into a medium-sized context length of 768 when NUC-encoded (Fig.1).

#### 2.2.2 K-mer Tokenization

With a vocabulary comprised of all nucleotide combinations of length *k*, K-mer Tokenization tokenizes the input as non-overlapping consecutive k-mers [25]. This reduces the sequence length by a factor of *k* and yields a vocabulary of size 4^*k*^. For a large *k*, the exponentially large vocabulary size introduces token sampling efficiency problems [23] and results in an explosion of parameters in the transformer’s embedding layer. This complicates the training and may not reflect the actual complexity of the data, as two k-mers are learned independently even when they are highly similar. Moreover, the model must learn and recognize each data signal in *k* alternative reading frames, taking up parameters/dimensions that would preferably be dedicated to other patterns.

#### 2.2.3 Byte Pair Encoding

BPE considers the most frequently occurring combinations of characters as tokens [34] and was first applied to nucleotide sequences in [23]. During training, BPE iteratively expands its vocabulary by merging the most frequent token pairs in the input corpus. This process is repeated until a prespecified vocabulary size (*vs*) is reached. During tokenization, BPE merges the subwords in the same order. BPE yields a highly efficient, fixed-size vocabulary of variable-length tokens. This method has three advantages over K-mer Tokenization: 1) a larger sequence length reduction for the same vocabulary size (Fig.1); 2) a higher token sampling efficiency; and 3) a better robustness against frameshifts. BPE has been shown to improve upon K-mer Tokenization in multiple NLM applications [23, 35], but still requires a large vocabulary to obtain strong compression [24].

#### 2.2.4 Convolutional Sequence Encoding

With CSE, nucleotide sequences are directly embedded into a high-dimensional space by means of a convolution (Fig.2). First, we convert an input sequence of length *l* to a 4 *× l* Position Weight Matrix (PWM), for example A = [1, 0, 0, 0]^*⊤*^ and N = [0.25, 0.25, 0.25, 0.25]^*⊤*^. A one-dimensional convolution layer (ReLU activated) with four input channels then transforms the PWM into *d*_*model*_ dimensions for the transformer using *d*_*model*_ learnable kernels. The stride is set to kernel size *k*, such that the sequence length is reduced *k* times, analogous to K-mer Tokenization. We add zero-padding to allow mini-batch training, and mask these positions during the attention operation.

**Figure 2:**
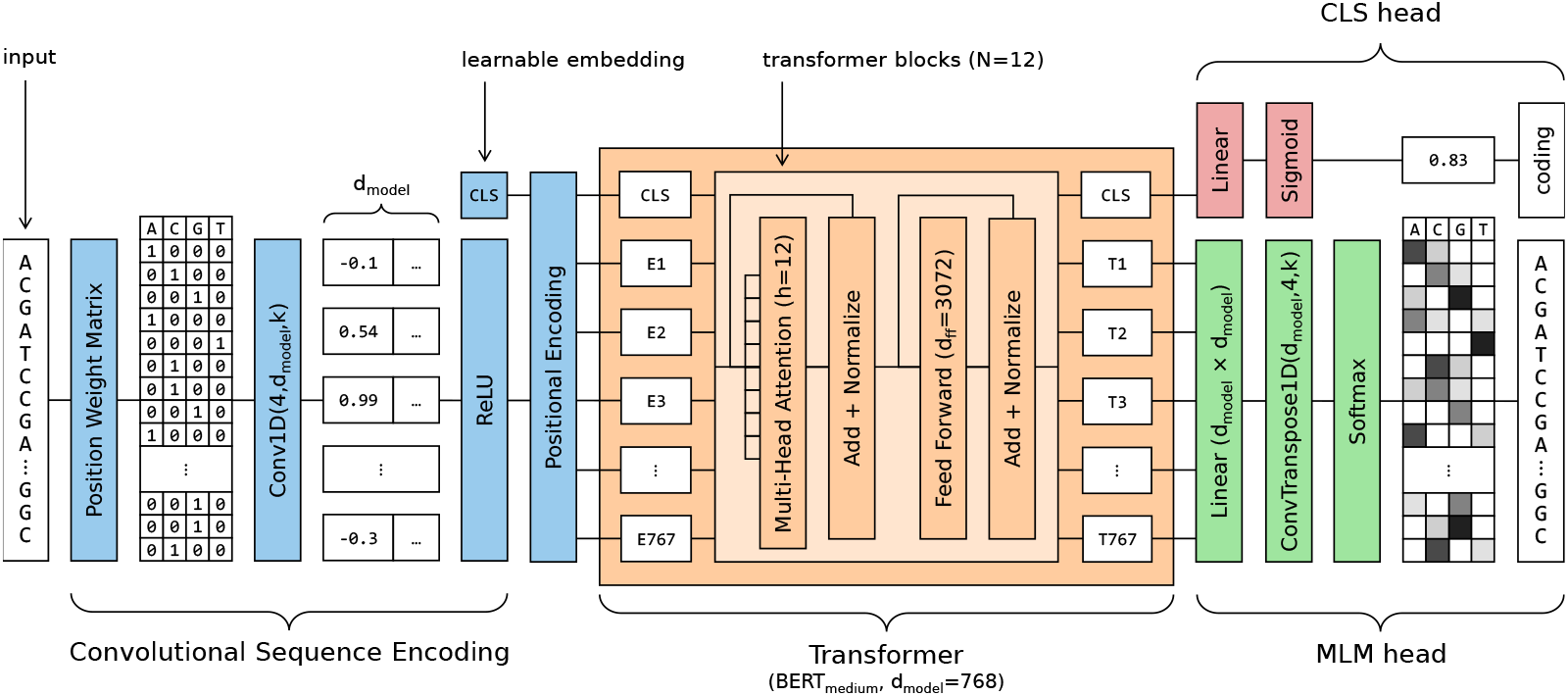
Architecture of lncRNA-BERT with Convolutional Sequence Encoding (CSE), adopting the BERT_medium_ transformer architecture [38]. CSE embeds a nucleotide sequence into *d*_*model*_ dimensions by means of a 1D convolution on its Position Weight Matrix, using *d*_*model*_ learnable kernels of size *k*. Fine-tuning tasks such as lncRNA classification are performed using an output head connected to the transformed CLS embedding. A dedicated MLM output head performs a transposed convolution, which enables masking and prediction at nucleotide resolution.

CSE effectively reduces the embedded sequence length with an efficient number of trainable parameters while maintaining nucleotide-level resolution. The method is inspired by the Vision Transformer [36], a similar approach for short DNA sequences was taken in [37]. Through the use of convolutions, CSE extracts important patterns from the data, seeing the input positions as combinations of nucleotides instead of independent tokens.

### 2.3 Language Model Architecture

We adapted the transformer encoder architecture as used by the BERT_medium_ [22, 38], with *N* = 12 transformer blocks, a dimensionality of *d*_*model*_ = 768, *d*_*ff*_ = 3072 nodes in the feed-forward layers, and *h* = 12 attention heads. lncRNA-BERT uses a medium-sized context length of *c* = 768 input positions. The transformed embedding of the CLS token was used as input to the lncRNA classification output head, which is a sigmoid-activated linear layer containing a single node. The model has 85M trainable parameters.

Our CSE-based models use a modified version of BERT (Fig.2). For classification, we replaced the CLS token with a learnable CLS embedding, as in ViT [36]. For the Masked Language Modeling (MLM) pre-training task, we enable nucleotide-level predictions by applying a softmax-activated, transposed convolution with stride/kernel size *k* on linearly transformed embeddings.

### 2.4 Training

We pre-train lncRNA-BERT for 7 days on an A100 MIG 4g.40GB GPU in the ALICE compute resources provided by Leiden University, after identifying an optimal model configuration based on pre-training, fine-tuning, and probing. The optimal model checkpoints were stored based on the validation set performance.

#### 2.4.1 Pre-training

MLM was used as a pre-training task, where tokens or nucleotides were selected with a probability of *p*_*mlm*_ = 0.15, of which 80% is masked and 10% is randomly replaced. The model’s task is to predict which tokens or nucleotides occur at the selected positions. With CSE, we use the IUPAC symbol ‘N’ to mask out nucleotides, as it indicates an equal probability for any of the four bases. The model is pre-trained using a cross entropy loss function, a batch size of 8, the Adam optimizer, and the learning rate schedule proposed in [22], with 32,000 warmup steps.

#### 2.4.2 Fine-tuning

LncRNA-BERT was fine-tuned for coding potential classification on 101,270 coding and 48,785 long noncoding RNAs from GENCODE and RefSeq. We optimized 100 epochs of 10,000 samples using Adam, a fixed learning rate of 10^*−*5^, and a batch size of 8. A binary cross entropy loss function was used, with the reciprocal class sizes as weights to counteract the class imbalance.

#### 2.4.3 Probing

Probing is used to evaluate the extent to which a pre-trained model encodes the coding potential in its sequence embeddings, without fine-tuning the weights of the network itself [25]. Hereto, we trained a small Multi-Layer Perceptron with a single hidden layer of 256 nodes on the mean-pooled output embeddings of a model. Optimization follows the same settings as during fine-tuning, except for an increased learning rate of 10^*−*4^.

### 2.5 Experimental Setup

We compared lncRNA-BERT to six published lncRNA classification algorithms. These include three classical approaches (CPAT [8], LncFinder [14], PredLnc-GFStack [13]), and three deep learning methods (feature-based LncADeep [16], sequence-based mRNN [17], and hybrid model RNAsamba [18]). LncADeep, RNAsamba, and mRNN are the best-performing, publicly available, human models in a recent benchmark [21]. Out-of-the-box models were used without re-training.

We experimented with two pre-training datasets (human, RNAcentral) and four encoding methods (NUC, K-mer, BPE, CSE) to determine an optimal lncRNA-BERT configuration. These comparisons were based on 500 pre-training epochs. The macro-averaged F1-score was used as the main performance metric, balancing precision and recall while equally weighing pcRNA/ncRNA to address class imbalance.

## 3 Results

### 3.1 LncRNA-BERT Outperforms and Matches State-Of-The-Art Performance in lncRNA Classification and Demonstrates a Better Generalizability

We selected 3-mer tokenization and CSE with *k* = 9 as the optimal encoding methods (Section 3.3), and benchmarked these configurations of lncRNA-BERT against six existing methods. As illustrated in Table 2, lncRNA-BERT matched state-of-the-art lncRNA classifiers in the two test sets and significantly outperformed all other methods in the most difficult RNAChallenge test set. This result proves the applicability of NLM for distinguishing coding from long non-coding RNA. In particular, we improved upon a previous sequence-based model (mRNN) by using a more advanced architecture and training procedure (BERT instead of RNN).

**Table 2:**
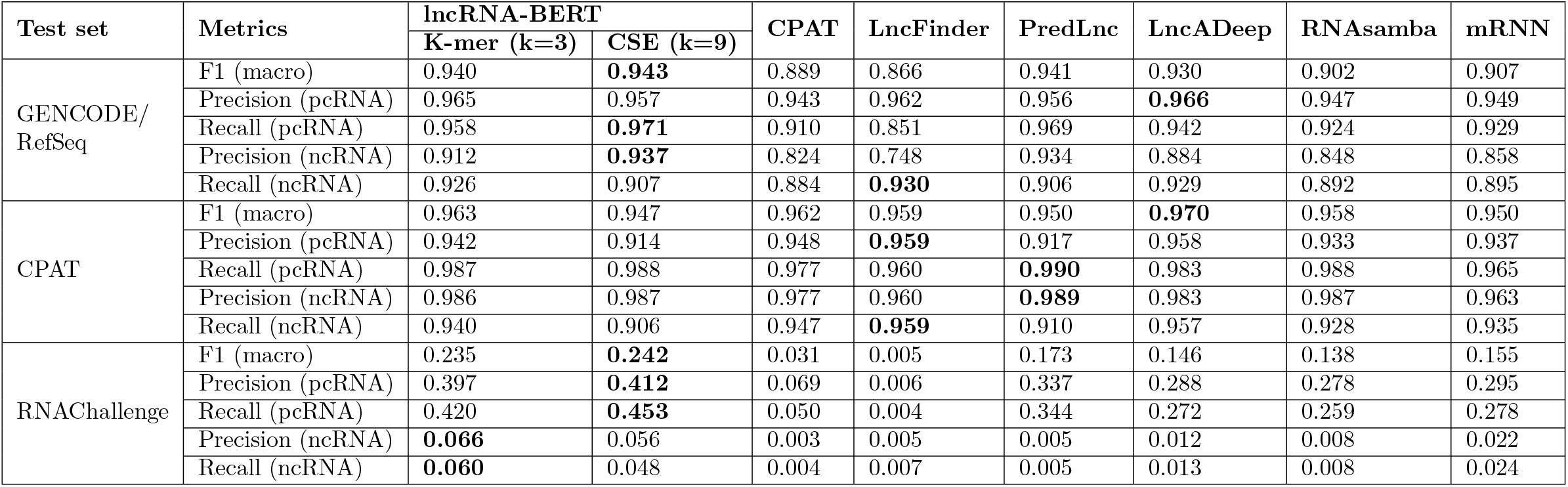
Comparison of lncRNA classifiers in 3 test sets.

Our BERT-based lncRNA classification method did not result in a large performance gain in comparison to the existing methods for two human test sets. Previous studies have set a high standard for lncRNA classification and a performance plateau is likely within reach (Section 4). In particular, for the CPAT test set, all methods performed almost equally well with a macro F1-score between 0.950 and 0.970 (lncRNA-BERT’s score is 0.963). The fact that the CPAT test set has been a well-known set for many years and contains more distinguishable pcRNAs and lncRNAs also makes true performance distinction difficult. Furthermore, some methods also benefit from sequence-extrinsic features, which cannot be learned by NLMs. For example, LncADeep performs alignments against a protein reference database to which lncRNA-BERT does not have access to. Nevertheless, we showed that coding potential is mostly a sequence-intrinsic characteristic, as lncRNA-BERT already distinguishes between pcRNA and lncRNA after self-supervised pre-training (Section 3.2).

Overfitting is a difficult problem in machine learning, which can cause some models to perform well on a particular test set but fail to generalize to unseen data. This potential bias in our analysis is mostly caused by the similarity between the training and test sets, which we mitigated for lncRNA-BERT by generating an independent train/test split after redundancy removal with CD-HIT (Section 2). The consistent performance of lncRNA-BERT on an independent test set shows proper generalization within human data competitive with established methods. The performance on the cross-species RNAChallenge benchmark also showed improvements in generalization across species compared to established methods.

### 3.2 Pre-Training on Human pcRNA/lncRNA Captures Coding Potential Without Access to Target Labels

The MLM pre-training task enabled our model to differentiate coding from long non-coding RNA without depending on target labels, given an appropriate pre-training dataset, as seen in the embedding spaces in Fig.3A. This affirms that the coding potential is a prominent sequence-intrinsic signal. Although lncRNA-BERT is not the first method to solely base its predictions on sequence patterns [7, 17, 39], it is the first to be capable of discriminating between pcRNA and lncRNA in a fully self-supervised manner. Fig.3B indicates that pre-training on human data leads to faster convergence and higher F1-scores (0.01-0.08 performance gain) compared to RNAcentral or no pre-training. This shows that pre-training on a proper dataset benefits classification.

**Figure 3:**
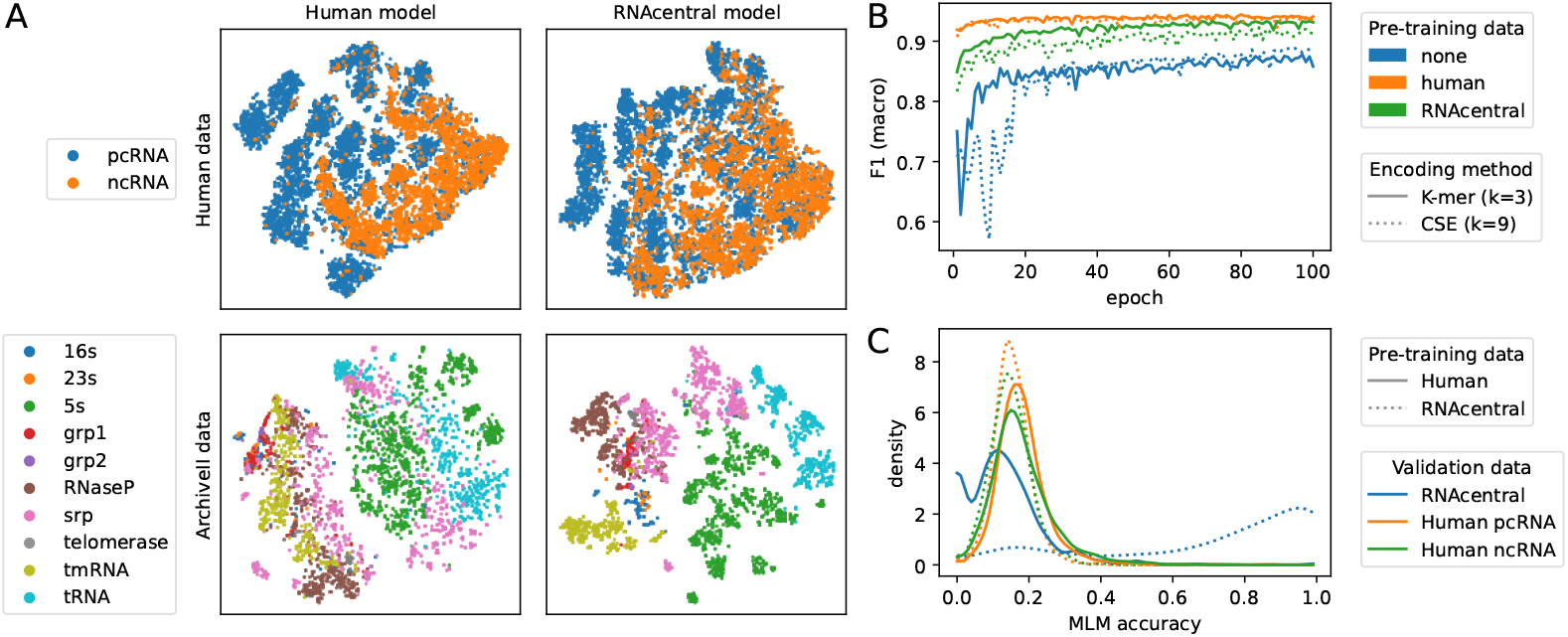
The effect of pre-training data on lncRNA-BERT. A: T-SNE visualizations of the embedding spaces of the human validation set and the ArchiveII dataset after pre-training lncRNA-BERT with 3-mer tokenization on human data or on multi-species data from RNAcentral. The human model better distinguishes pcRNA from ncRNA than the RNAcentral model, while the latter generates an improved separation between structural families in ArchiveII. B: Learning curve of the macro-averaged F1-score during 100 training epochs on the lncRNA classification task, for different pre-training configurations and encoding methods. The models benefit from pre-training, specifically on human data, leading to faster convergence and increased performance. C: Density plot of the MLM accuracy per sequence (% correctly predicted tokens) for lncRNA-BERT with 3-mer tokenization when pre-trained/evaluated on human pcRNA/ncRNA data or cross-species ncRNA from RNAcentral. The RNAcentral model achieves a high MLM accuracy (mean: 70%) on the RNAcentral validation set but performs worse than the human model on lncRNA from the human validation set (mean: 15% versus 18%). The models slightly favor pcRNA over ncRNA.

Pre-training on RNAcentral biases the model towards ncRNA types other than human pcRNA and lncRNA. This is reflected in the per-sequence MLM accuracy (% of correctly predicted tokens) in Fig.3C, which shows that the RNAcentral model performs well on sequences from RNAcentral (mean: 70%), but does not achieve the same accuracy on human lncRNAs (mean: 15%). The same can be concluded from the embedding spaces generated by both models. The RNAcentral model embeds pcRNA and ncRNA less distinctly than the human model. We also generated embedding spaces for the ArchiveII dataset, containing multi-species ncRNAs from 10 structural families [40]. Here, the RNAcentral model successfully separates the different classes, in contrast to the human model. Hence, the patterns learned from RNAcentral do not generalize well to human pcRNA/lncRNA, and vice versa. This may be caused by the abundance of specific ncRNAs in RNAcentral, as well as the complete lack of pcRNA. Moreover, lncRNAs are less well-conserved across evolution [12], complicating the detection of coding potential in a cross-species dataset.

Our results reveal a potential weakness of existing RNA language models, which use RNAcentral as the main data source, making them less suitable for pcRNA/lncRNA classification.

### 3.3 Convolutional Sequence Encoding Improves Pre-Training on Longer RNA Sequences

We pre-trained, probed, and fine-tuned BERT models with different encoding methods (Fig.4), and identified CSE (*k* = 9) and 3-mer tokenization as the most suitable for coding potential classification.

**Figure 4:**
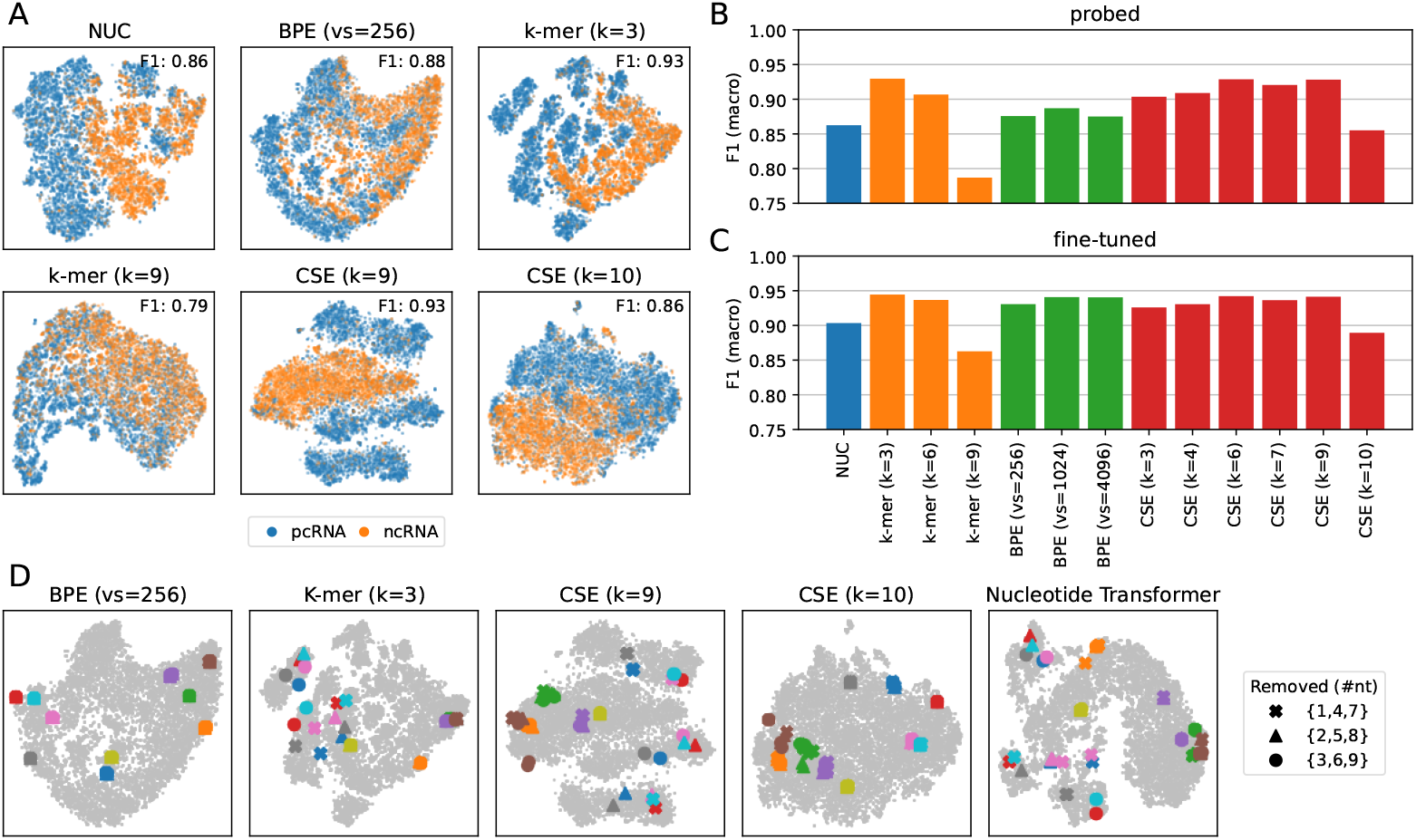
Embedding spaces and performance of lncRNA-BERT for different encoding methods. Mean pooling is used to retrieve sequence-level embeddings. A: T-SNE visualizations of the embedding spaces of the validation set after pre-training models with different encoding methods. K-mer Tokenization with *k* = 3 and CSE with *k* = 9 lead to the best distinction between pcRNA and lncRNA, reflected in the achieved F1-score when probed (0.93). CSE improves upon K-mer for *k* = 9. B-C: Macro-averaged F1-scores on the lncRNA classification task for different encoding methods, using probing (B) or finetuning (C). K-mer Tokenization with *k* = 3 leads to the highest performance for both probing (0.93) and fine-tuning (0.94). D: T-SNE visualizations of the embeddings of models with different encoding methods, which change depending on the reading frame of the input. Assessed by removing up to 9 nucleotides from the start of 10 randomly selected sequences (indicated by color) in the validation set. The sequence-level embedding does not change for BPE, and CSE when *k* = 10. Sequence embeddings jump across three different coordinates when using K-mer Tokenization or CSE with *k* divisible by 3.

Of all encoding methods that achieve a large sequence length reduction (6 ≥ times), CSE leads to the most effective models. This is reflected in the embedding spaces (Fig.4A) as well as in the obtained F1-scores after probing (Fig.4B). Specifically, CSE and K-mer Tokenization obtain a probing F1-score of 0.93 versus 0.91 for *k* = 6, and 0.93 versus 0.79 for *k* = 9, respectively, on the validation dataset. CSE’s fine-tuning scores are also higher for these configurations (Fig.3C). The reduced performance of K-mer Tokenization can be explained by the limitations of using a large token vocabulary (e.g. 4^9^ for 9-mers), which include reduced sampling efficiency [23] and a high embedding layer parameter count (e.g., 4^9^ *× d*_*model*_ for 9-mers). By contrast, CSE sees k-mers as combinations of nucleotides and maintains a fixed number of parameters (4 *× d*_*model*_ *× k*) by learning important patterns from the data. BPE encodes sequences into fully independent tokens, such as the k-mer approach, contributing to why it is outperformed by CSE for *υs* ∈ [256, 1024, 4096] when probed.

Although superior for long sequences, CSE is outperformed by K-mer and BPE when the token size is small and fine-tuning is allowed. In Fig.4C, CSE with *k* = 3 results in a fine-tuning F1-score of 0.93, while the 3-mer and BPE for *vs* = 256 and *vs* = 1024, achieve 0.94, 0.93, and 0.94 respectively. Shorter tokens enable attention at a higher resolution, thereby improving contextualized embeddings. Seeing input positions as predefined, independent tokens also helps the model discriminate between sequences. In comparison, CSE must learn important patterns before BERT can condition on them, making the model less stable than when using a tokenizer. Issues specific to tokenizers, such as sampling efficiency and parameter count blow-ups, are less prevalent for smaller vocabularies. This specifically applies to finetuning, during which the model can prioritize important tokens over less important tokens. Consequently, BPE and K-mer tokenization show a larger performance gain from probing to fine-tuning compared to CSE.

Another factor that affects classification performance is the sequence length coverage of each encoding method (Fig.1). A longer coverage allows the model to consider a larger part of the sequence in its predictions, explaining why 3-mer tokenization is superior to NUC, as shown in Fig.4. An extended context length may also hinder the model from learning important local signals such as CSE *k* = 10. The context length of a model with 3-mer tokenization (3 *×* 768 = 2304) was shown to be sufficient for classifying coding potential. The biological implication here is that the coding potential of long RNA transcripts is usually inferrable from the first 2304 nucleotides. This, in combination with its small vocabulary, explains why a model with 3-mer tokenization achieves a similar fine-tuning performance to CSE with *k* = 9 (0.944 versus 0.941), despite having a smaller context length.

### 3.4 Three-Base Periodicity in Sequence Encoding Method Benefits Performance and Affects Embedding Space

Encoding methods that align with the three-base periodicity of coding RNA have been shown to better distinguish pcRNAs from lncRNAs, owing to their sensitivity to biological reading frames. For example, CSE with *k* ∈ [6, 9] improves upon *k* ∈ [7, 10]. However, there is also a negative effect of the reading frame sensitivity of the three-base periodic encoding methods as illustrated in Fig.4D, in which the same pcRNA sequence could jump between the three groups when observed in different reading frames. While input position embeddings are reading frame dependent (particularly for K-mer and CSE), the aggregated (mean) embedding at the sequence level should remain the same, as the sequence’s meaning is preserved. Fig.4D shows that this applies to BPE or CSE with *k* = 10, but not for K-mer or CSE with *k* as a multiple of 3. Moreover, the figure shows that the Nucleotide Transformer, which uses 6-mer tokenization, suffers from the same issue. This effect was not observed in the non-coding sequences.

These observations demonstrate how biological reading frames in the data affect the behavior of three-base periodic models, which appear to favor in-frame signals. These signals are easier to learn, because their periodic organization aligns with the input positions of the model. For example, from ‘ACT TGA ACT’ it is easier to learn that ‘TGA’ follows ‘ACT’, compared to learning that ‘GAA’ follows ‘CTT’. We expect the presence/absence of such easily recognizable signals to cause the observed shifts between the embeddings in Fig.4D. Three-base periodicity is less evident in lncRNA, causing a lower MLM accuracy (Fig.3) but a better embedding consistency than pcRNA. The frameshift sensitivity is mitigated by setting *k* to a value not divisible by 3, which breaks up the three-base periodicity.

### 3.5 Embedding Spaces of pcRNA/lncRNA Data Reveal Differences Between NLMs

We compared lncRNA-BERT to previously released NLMs by visualizing the embedding spaces of the validation set in Fig.5. lncRNA-BERT is pre-trained on human pcRNA and lncRNA, in contrast to other RNA NLMs, which use RNAcentral as the main data source. Excluded from Fig.5 are ERNIE-RNA, RNABERT, RNAErnie, and RNA-FM [26–29]. These methods were only trained on non-coding RNAcentral data, making them less suitable for inference on pcRNAs and lncRNAs (Fig.3). BiRNA-BERT and RiNALMo also utilize the RNAcentral dataset, but augment it with data from RefSeq [41] and Rfam/Ensembl [30]. Nevertheless, the embedding space of BiRNA-BERT did not separate pcRNA from ncRNA as clearly as lncRNA-BERT. Only the RiNALMo model (650M parameters) generated a clearer distinction, but was 7.6 *×* larger than lncRNA-BERT (85M parameters) and was pre-trained with a limited context length (*<* 1024 nt).

**Figure 5:**
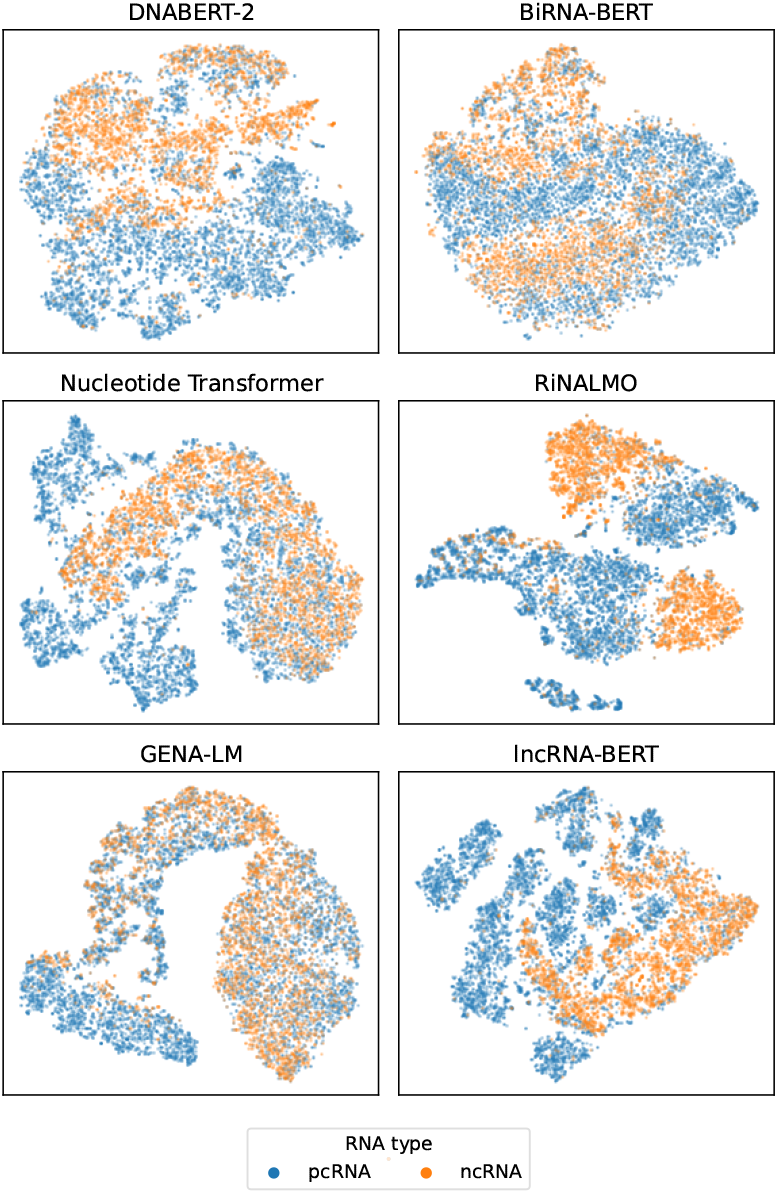
T-SNE visualizations of the embedding spaces of the validation set, generated by different NLMs. RiNALMo generates the clearest distinction between coding and non-coding transcripts, followed by lncRNA-BERT and DNABERT-2. BiRNA-BERT, Nucleotide Transformer (v2, multi-species), and GENA-LM (base, T2T) differentiate the two types of RNA less clearly.

Fig.5 shows the competitiveness of lncRNA-BERT with other NLMs on this dataset and task, while also being the most parameter-efficient. Interestingly, DNA NLMs such as DNABERT-2 and Nucleotide Transformer (v2) show some potential for RNA classification. This indicates that DNA models may be generalized to RNA tasks, although not as well as to RNA-specific models.

## 4 Discussion

The human RNA language model proposed in this work, lncRNA-BERT, demonstrates state-of-the-art performance in classifying RNAs as coding or long non-coding (Table2). Pre-training on human pcRNA/lncRNA from GENCODE, RefSeq, and NONCODE leads to a clear distinction between the two classes, showing that an NLM can learn coding potential from sequence data without relying on target labels (Fig.3A). LncRNA-BERT stands apart from other RNA language models through its longer context length, smaller model size, and the use of human pre-training data, which enhances classification performance compared to models pre-trained with RNAcentral (Fig.3B).

A performance plateau may have been reached for binary coding potential classification, which could explain why lncRNA-BERT did not consistently outperform existing methods. Even a simple logistic regression algorithm (CPAT) can reach an F1-score of 0.89 on GENCODE/RefSeq (Fig.2), indicating the triviality of this task. At the same time, the problem is complicated by the false assumption of an unambiguous separation between pcRNAs and lncRNAs. Some lncRNAs are known to contain short ORFs that encode functional micro-peptides [10], whereas some pcRNAs can have lncRNA-like regulatory functions [11, 12]. Hence, it is impossible to obtain 100% accuracy without overfitting to the human annotation system. Here, a method such as LncADeep benefits from incorporating protein alignment data in its predictions, which is also dependent on currently available knowledge. In contrast, lncRNA-BERT distinguishes coding from non-coding RNA without access to annotations, paving the way for a more nuanced view of the two RNA classes in future work.

We demonstrated that the choice of encoding method and pre-training data is critical for lncRNA-BERT’s performance. This highlights two key areas for potential improvements in NLMs in future studies. Regarding encoding methods, each have a different effect on the obtained sequence coverage, resolution, efficiency, and information retention. NUC represents nucleotides individually rather than combining them into single input positions, resulting in a demanding attention operation and a limited context length. While context length may be extended through architectural advancements, such as Attention with Linear Biases [23, 41] and Flash Attention [23, 30], adopting a more compressive encoding method is still advisable. BPE reduces the sequence length more efficiently than NUC and K-mer, but leads to reduced probing performance. This indicates that BPE models are less capable of recognizing coding potential after pre-training, in comparison to using K-mer or CSE. We attribute this to BPE’s frequency-based vocabulary, which may not be biologically relevant and breaks up the three-base periodicity. Inconsistent token size also introduces problems for other fine-tuning tasks that require nucleotide resolution. Of all encoding methods, CSE enables the strongest compression by using learnable convolutions to recognize complex patterns from RNA sequences. Despite this, 3-mer tokenization obtains similar performance, indicating that a context length of 2,304 nucleotides is sufficient for recognizing the coding potential of most RNAs. lncRNA-BERT with CSE may potentially lead to superior performance when fine-tuned for other tasks that require longer context lengths. Ultimately, the choice of encoding method depends on the characteristics of a specific problem and dataset, and identifying a universal solution requires further evaluation on diverse fine-tuning tasks.

Regarding pre-training data, an NLM requires human pcRNA/lncRNA to achieve maximum performance in distinguishing the two classes, as shown in Fig.3 and Fig.5. This demonstrates the necessity of a sufficient amount of task-specific pre-training data. Nevertheless, we anticipate that lncRNA-BERT and other RNA language models may be improved by increasing the dataset size when a proper class balance and sequence diversity are ensured. Adding genetic variation has helped DNA language models generalize over subtle signals between different individuals and phylogenetic signals between species. For example, pre-training the Nucleotide Transformer with data from the 1000 Genomes Project enables it to perform variant prioritization [25], and GENA-LM can be used for taxonomic classification after pre-training on multi-species data [24]. In contrast, our results show that using the 37M multi-species RNAcentral dataset as the only pre-training source does not improve lncRNA-BERT, but results in a bias towards RNA types other than pcRNA and lncRNA (Fig.3C). Algorithms such as CD-HIT and MMSeqs2 can remove redundant sequences from datasets like RNAcentral, which may be the key to ensuring proper data balance and diversity. RiNALMo implements this approach, but excludes pcRNA [30]. Hence, pretraining a model with an efficient encoding method on a well-balanced dataset encompassing all types of RNA may yield the next generation of RNA NLMs.

## Data Availability Statement

The data supporting the findings of this study are openly available from GitHub at https://github.com/luukromeijn/lncRNA-Py, GENCODE (v46) at https://www.gencodegenes.org/, RefSeq (v225) at https://www.ncbi.nlm.nih.gov/refseq/, NONCODE (v6) at http://v6.noncode.org/, and RNA-central (v24) at https://rnacentral.org/.

